# Prenatal maternal infection promotes maternal microchimeric cells to alter infection risk in male offspring

**DOI:** 10.1101/2025.01.17.633596

**Authors:** Brittany Jimena, Daniel Kazimierczyk, Simon Kazimierczyk, Haley Moya, Eva Shin, Li Li, Parimal Korgaonkar, Caryn Porter, Brian Seed, Bobby J. Cherayil, Nitya Jain

## Abstract

Vertically transferred maternal cells or maternal microchimeric cells (MMCs) engraft the fetus and persist in offspring for long periods of time. How altered maternal immune states arising from infection affect MMCs and their function in offspring is poorly understood. Here, we show that pregnancy-associated transient maternal infection alters MMCs to differentially regulate immunity in offspring. In male offspring of dams previously infected with *Yersinia pseudotuberculosis*, MMCs confer a pro-inflammatory type 17 T effector phenotype that leads to enhanced protective immunity to an unrelated *Salmonella* infection. Thus, acquired maternal cells imprinted by microbial exposure during pregnancy exert an antigen agnostic and sex-differential effect on offspring immunity, and may potentially be targeted to deliver immune benefits to infants in the vulnerable early life period.

## Introduction

Exposure to infectious agents during pregnancy can impact offspring immune responses to infections and vaccinations(1). Chronic infection during pregnancy can affect infant health independently of pathogen transmission. For instance, HIV-exposed but uninfected infants have reduced levels of maternal antibodies and show increased susceptibility to infection by unrelated pathogens compared with infants not exposed to HIV(2). Human malarial and helminth infection during pregnancy correlates with impaired IgG responses to *Haemophilus influenzae* and diphtheria antigens(3). Vertically transferred maternal immunity including antibodies and cytokines have a large influence on immune responses in early life. Maternal cells that engraft the fetus during gestation, called maternal microchimeric cells (MMCs), also have the potential to influence offspring immunity(4).

Acquired MMCs are retained in offspring for long periods of time and are proposed to have both harmful and beneficial roles(5, 6). Alloimmune inflammatory responses targeting genetically disparate MMCs present in heart tissues contribute to the pathogenesis of cardiac injury in neonatal lupus(7). Acquired maternal T cells initiate islet inflammation and predispose to diabetes in an animal model of autoimmunity(8). MMCs contribute to tissue repair(9, 10), and substitute for missing fetal immune components in immune-deficient animals(11, 12). MMCs also regulate offspring immunity through antigen-specific and non-specific mechanisms. Immunodeficient infants with Epstein-Barr virus (EBV) and CMV infection harbor maternal CD8^+^ T cells that are reactive to EBV and CMV antigens respectively (13, 14). In an animal model of preconceptual infection with OVA expressing *Listeria*, OVA-specific maternal T cells are transferred to the fetus during pregnancy(15). The presence of these chimeric T cells in offspring correlates with less severe disease following infectious challenge in early life. In addition to contributing to pathogen-specific immunity in offspring, MMCs prime the expansion of fetal Treg cells that avert maternal-fetal conflict during pregnancy(16) and confer reproductive fitness in female offspring(17). MMCs in the fetal bone marrow promote the differentiation of CD11b^+^GR-1^lo/neg^ monocytes that corelate with protection from neonatal CMV infection in mice(18). MMCs are also associated with reduced risk of early life infections in humans, especially in infant boys(18).

MMCs may have sex-differential functions in offspring, contributing to protection against complications in next-generation pregnancies of female offspring(17), while promoting immunity to early life respiratory infection in male offspring(18). Offspring female sex is also associated with increased presence of MMCs in infants and young children(19, 20). Maternal inflammation during pregnancy is particularly associated with sex-differential effects on offspring. Male rat blastocysts are more susceptible to pregnancy associated inflammation than female blastocysts(21). Maternal immune activation during pregnancy with poly I:C leads to social and behavioral changes specifically in male offspring(22). Whether maternal infection influences MMCs and their function is unknown.

Most infections experienced during human pregnancies are mild and often self-resolve(23). In this study, we leveraged a pregnancy-associated, low-pathogenicity transient infection model to determine the impact of maternal immune activation on MMCs in offspring. Male offspring of previously infected dams harbored more MMCs that promoted the development of type 17 T effector cells and contributed to better health outcomes after infection as young adults. Our studies identify a sex-specific function of MMCs that is revealed upon maternal infection.

## Results

### Pregnancy- associated maternal infection increases MMCs in offspring

To identify chimeric cells in mouse offspring, we generated female heterozygous PhAM^excised^ mice (Dendra^HET^; fully backcrossed to C57BL/6 strain background) that express a green fluorescent protein called Dendra from a ubiquitous mitochondrial promoter(24). Allogeneic matings, that have previously been shown to increase maternal cell trafficking to offspring (25), of Dendra^HET^ female with Balb/c male mice were set up to generate WT progeny (henceforth called Dendra^WT^ to indicate WT progeny of Dendra-expressing mothers) (**Fig. 1A**). Dendra^WT^ progeny (that do not express endogenous Dendra-g protein) were analyzed for the presence of Dendra-g^+^ green-fluorescent maternal cells by flow cytometry. To assess the impact of pregnancy-associated infection on MMCs, we utilized a published animal model of transient, low-pathogenicity intestinal infection with an attenuated *yopM* mutant strain of *Yersinia pseudotuberculosis* (26, 27). Timed-pregnant Dendra^HET^ dams were infected with *yopM* mutant at E7.5 when primitive hematopoiesis is initiated in the yolk sac (YS). *yopM* infected dams experienced a transient weight loss after infection with peak bacterial shedding in stools at 3-4 days post infection (**Supp Fig 1A, B**). Maternal *yopM* infection altered splenic distribution of numerous immune subtypes, with a drop in total CD45^+^ hematopoietic cells 5 days after infection compared to uninfected pregnant dams (**Supp Fig 1C, D**). Within CD45^+^ cells, there were significant redistribution of immune subtypes with increases in CD11b^+^ myeloid cells including CD11b^+^F4/80^+^ macrophages and CD11b^+^Ly6G^+^Ly6C^+/neg^ cells (**Supp Fig 1C, D**).

**Figure 1:**
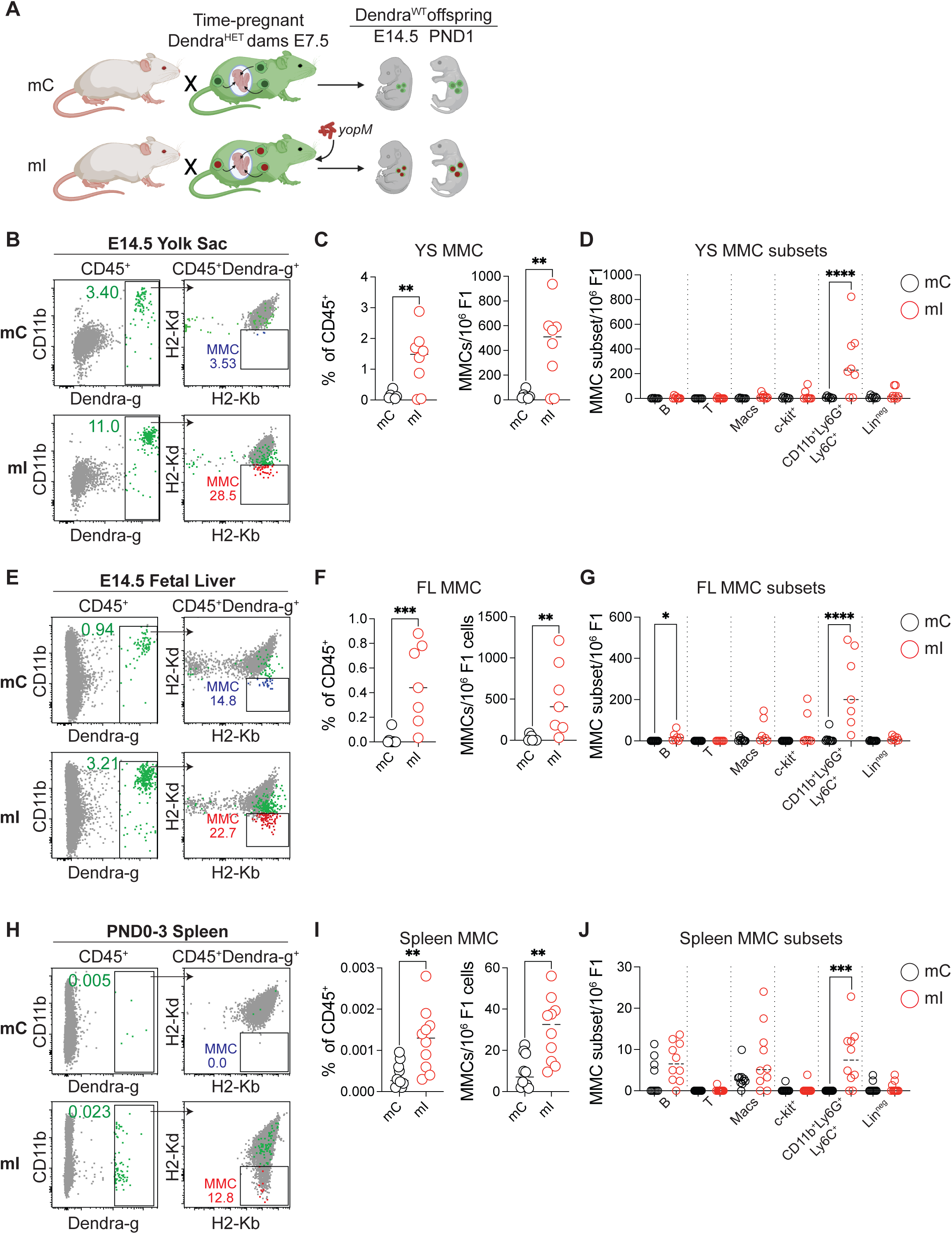
Pregnancy-associated maternal infection increases MMCs in offspring. (A) Allogeneic matings of Dendra^HET^ females with BALB/cJ males were set up to generate Dendra^WT^ offspring. Timed-pregnant Dendra^HET^ dams were orally administered *Y. pseudotuberculosis yopM* at embryonic day 7.5 (E7.5). Dendra^WT^ offspring of infected (mI) and control (mC) mothers were analyzed at E14.5 and postnatal day 1 (PND1). (B) E14.5 Yolk Sac (YS): (*Left*) Representative flow cytometry dot plots showing gating strategy to identify Dendra-g^+^ cells among CD45^+^ cells. Numbers indicate frequency of Dendra-g^+^ cells within CD45^+^ gate. (*Right*) Dendra-g^+^ cells were further analyzed for expression of H2-Kb and H2-Kd and MMCs were identified as Dendra-g^+^H2-Kb^+^H2-Kd^neg^. Numbers indicate frequency of MMCs within Dendra-g^+^ gate. (Gray: Total CD45^+^ cells; Green: Dendra-g^+^ cells; Blue: MMCs in mC offspring; Red: MMC in mI offspring) (C) E14.5 YS: (*Left*) Frequency of MMCs within CD45^+^ cells. (*Right*) Numbers of MMCs per million F1 cells. (mC n=7, mI n=8). (D) E14.5 YS: Subsets of MMCs were identified as CD19^+^ B cells, CD3e^+^ T cells, CD11b^+^F4/80^+^ macrophages (macs), c-kit^+^ cells, CD11b^+^F4/80^neg^Ly6G^+^Ly6C^+^ myeloid cells and Lin^neg^ (Lin: CD3e, CD19, CD11b, F4/80, Ly6G, Ly6C, c-kit). Plot shows number of MMCs per million F1 cells. (mC n=7, mI n=8). (E) E14.5 Fetal Liver (FL): Gating strategy to identify MMCs as in (**B**). (F) E14.5 FL: Frequency and number of MMCs as in (**C**) (mC n=9, mI n=7). (G) E14.5 FL: MMC subsets as in (**D**) (mC n=9, mI n=7). Data in **B-G** are representative of 2 independent experiments using 2 pregnant dams per group and 3-5 offspring per pregnancy. (H) PND0-3 Spleen: Gating strategy to identify MMCs as in (**B**). (I) PND0-3 Spleen: Frequency and number of MMCs as in (B) (mC n=10, mI n=10). (J) PND0-3 Spleen: MMC subsets as in (C) (mC n=10, mI n=10). Data in **H-J** are from 5 independent experiments using 3-5 pregnant dams per group and 2-5 offspring per pregnancy. **C, F, I**: t-test ** < 0.0021, *** <0.0002. **D, G, J**: One-way ANOVA Adjusted P-value: *** <0.0002, **** <0.0001. Summary of statistical analyses are shown in **Supplemental Data 1**.

We first compared the distribution and phenotype of MMCs in the extraembryonic YS and fetal liver (FL) tissues of concepti from naïve (mC) versus infected (mI) dams at E12.5-E14.5. Littermate microchimeric cells that are acquired from adjacent fetuses in multi-embryonic murine pregnancies(28) were excluded from analyses by staining with MHC haplotype antibodies to identify cells of maternal B6 origin (**Fig. 1 B, E, H**). Maternal infection during pregnancy increased the frequency and numbers of MMCs in both the YS and FL (**Fig. 1B-G**). As reported previously(18), MMCs were a heterogeneous mix of CD45^+^ hematopoietic cells including innate and adaptive immune cells (**Fig. 1D, G, Supp Fig. 2B-E**). However, maternal infection specifically increased the frequency of CD11b^+^ MMCs in the YS and FL of fetuses (**Fig. 1D, G**). The CD11b^+^ cells were primarily F4/80^neg^Ly6G^+^Ly6C^+^, a phenotype that resembled murine granulocytic myeloid derived suppressor cells (MDSCs) (**Supp Fig. 2B, D**)(29). We also analyzed newborn mice (PND0-3), and while the overall frequency of MMCs in the spleen was significantly lower compared to fetal tissues, there were similar increases in the frequency and numbers of MMCs and CD11b^+^ MMCs in pups delivered by previously infected dams compared to naïve dams (**Fig. 1H-J**).

**Figure 2:**
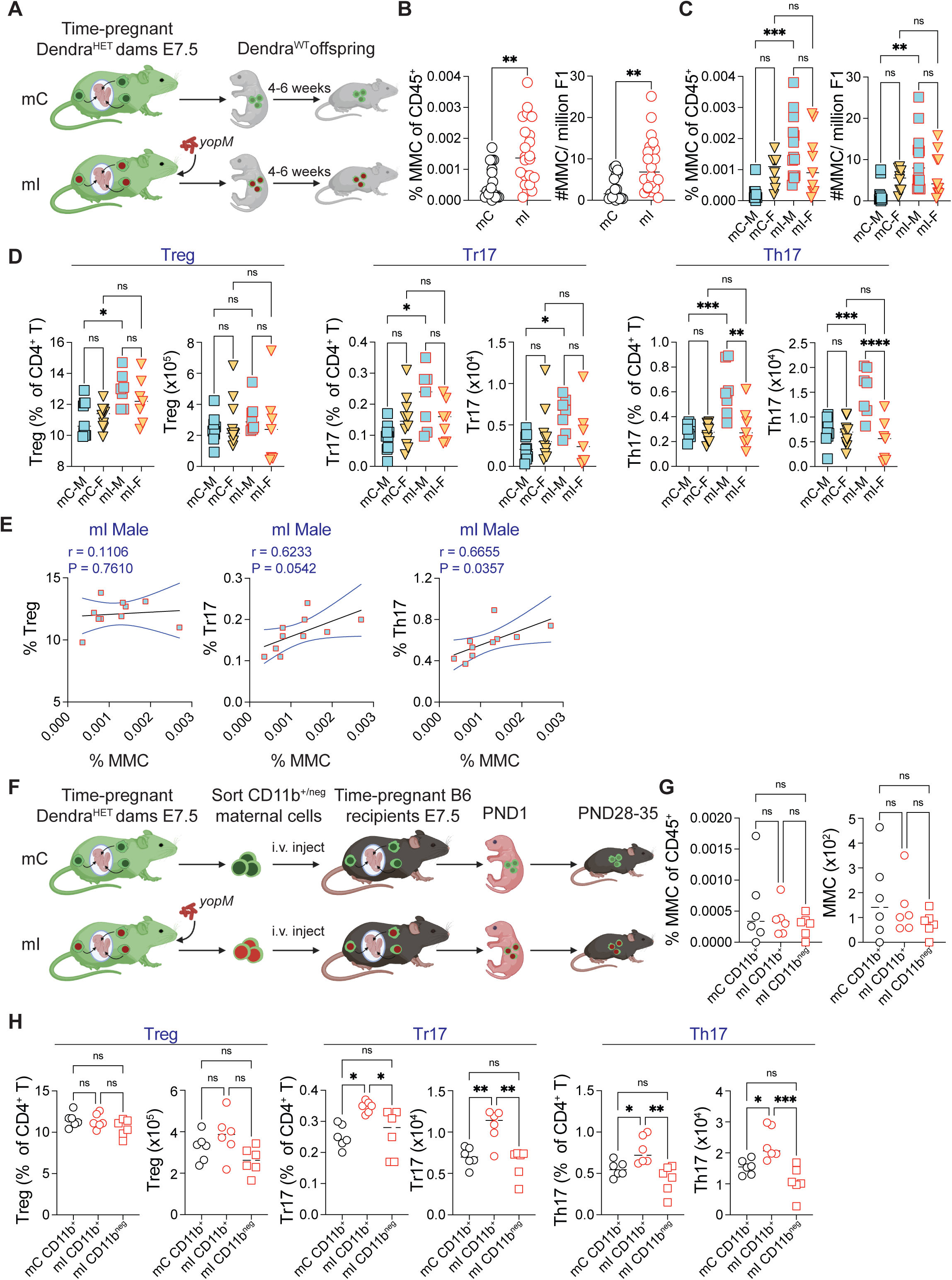
Pregnancy-associated maternal infection increases MMCs in male offspring that correlates with increased splenic CD4^+^ RORγt^+^ T cells. (A) Dendra^WT^ offspring of uninfected (mC) or previously infected dams (mI) were analyzed at 4- 6 weeks of age. (B) Frequency and numbers of MMCs within CD45^+^ cells in spleen (mC n=17, mI n=22; t-test P ** < 0.0021). (C) Frequency and numbers of MMCs within CD45^+^ cells in spleens of male and female mice. (mC male n=9, mC female n=8, mI male n=13, mI female n=9; data are from 5 independent experiments using 2-3 pregnant dams per group and 2-5 offspring per pregnancy). (D) Frequency and numbers of CD4^+^FOXP3^+^ Treg cells, CD4^+^FOXP3^+^RORγt^+^ Tr17 cells and CD4^+^RORγt^+^ Th17 cells in spleen. (mC male n=9, mC female n=10, mI male n=7, mI female n=7; data are from 3 independent experiments using 2-3 pregnant dams per group and 2-4 offspring per pregnancy). (E) Correlation analysis of the percentage of splenic MMCs versus the percentage of Treg, Tr17 and Th17 cells in male mI Dendra^WT^ offspring. The black line represents the linear regression, and the blue line indicates the nonparametric Spearman correlation with a 95% confidence interval. (F) Timed-pregnant Dendra^HET^ dams at post-coital day 7.5 were orally administered *Y. pseudotuberculosis yopM*. Five days post-infection, MACS enriched CD11b^+^ and CD11b^neg^ cells from spleens of uninfected (mC) and infected (mI) dams were i.v. injected into timed-pregnant B6 female recipients at post-coital day 7.5. Offspring of recipient B6 dams were analyzed at PND28- 35. (G) Frequency and numbers of Dendra-g^+^ MMCs within CD45^+^ cells. (mC CD11b^+^ n=6, mI CD11b^+^ n=6, mI CD11b^neg^ n=6) (H) Frequency and numbers of Treg cells, Tr17 cells and Th17 cells in spleen. (mC CD11b^+^ n=6, mI CD11b^+^ n=6, mI CD11b^neg^ n=6). Data in **F-H** are representative of 2 independent experiments using 2 pregnant dams per group and 4-6 offspring per pregnancy. One-way ANOVA adjusted P-value: ‘ns’ not significant, * <0.033, ** < 0.0021, *** <0.0002, **** <0.0001. Summary of statistical analyses are shown in **Supplemental Data 1**.

### Increased persistence of MMCs in male offspring of previously infected dams

We next assessed whether pregnancy-associated maternal infection altered persistence of MMCs in adult offspring. We analyzed mice after weaning at 4-6 weeks of age (**Fig. 2A**) and consistent with the early time points, observed an increase in frequency and numbers of MMCs in the spleen of adult offspring of previously infected dams (**Fig. 2B**). A secondary analysis to address sex as a biological variable revealed that male offspring of previously infected mothers harbored significantly more MMCs compared to male offspring of control mothers (**Fig. 2C**), that was confirmed by a two-way ANOVA analysis that yielded a significant sex and maternal infection interaction (**Supp Fig. 3A**)(30). A reanalysis of the fetal data segregated by sex, however, did not show significant interaction even though male fetuses appeared to have increased MMCs in the YS and FL during gestation (**Supp Fig. 3B, C**). These data suggested that postnatal mechanisms likely enforce sex-specific differences in MMC homeostasis.

**Figure 3:**
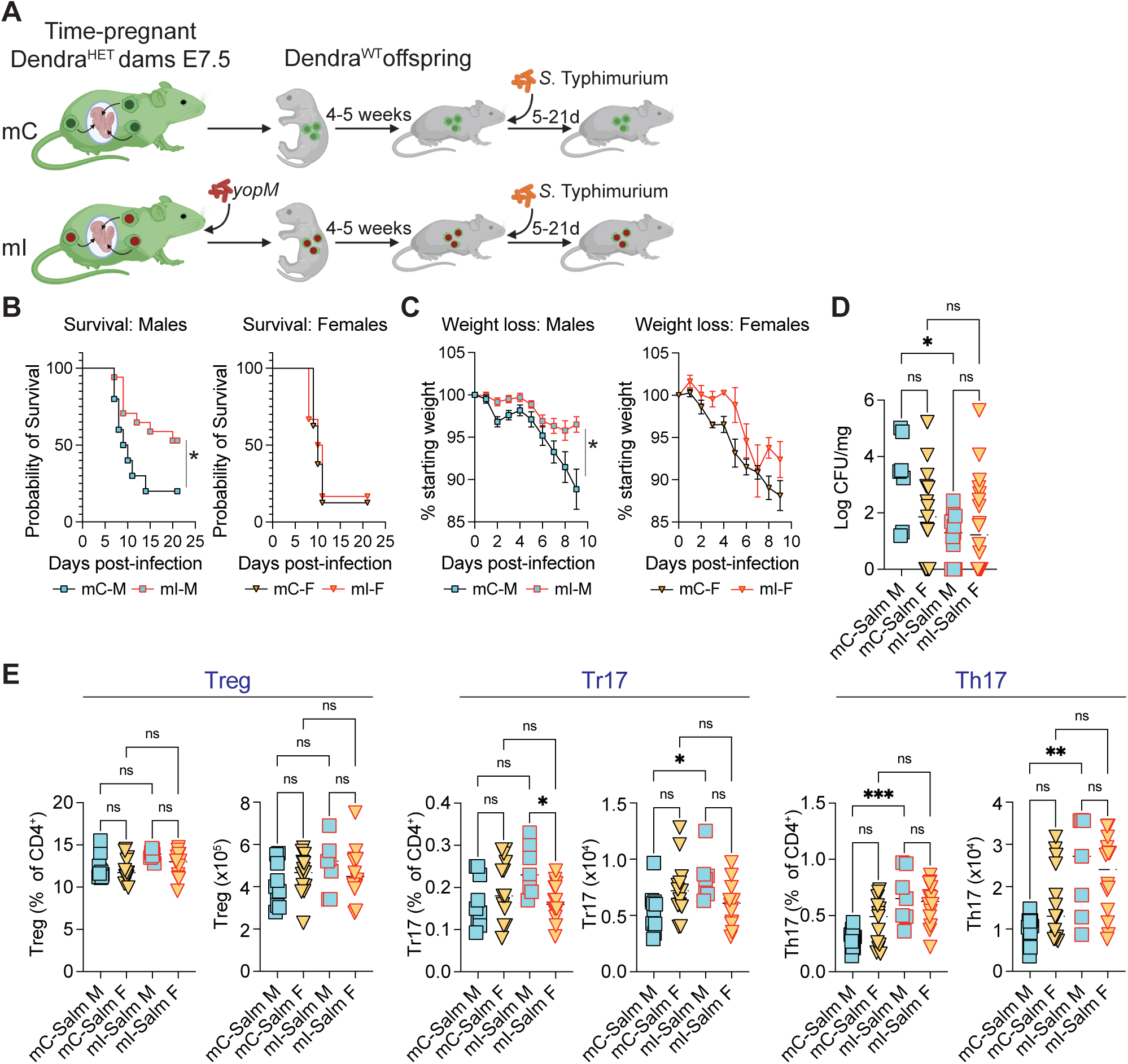
Pregnancy-associated maternal infection alters susceptibility of male offspring to Salmonella infection. (A) Dendra^WT^ offspring of uninfected (mC) or previously infected dams (mI) were orally administered *Salmonella* Typhimurium (SL1344 mutant) at PND28-35. (B) Survival curve of male and female offspring after *Salmonella* infection. (mC-Salm M n=10, mI-Salm M n=17, mC-Salm F n=8, mI-Salm F n=6; P * <0.03) (C) Percent weight change after *Salmonella* infection. (mC-Salm M n=9, mI-Salm M n=17, mC- Salm F n=6, mI-Salm F n=5; P * <0.03). (D) Splenic bacterial burden 5 days after *Salmonella* infection. (mC-Salm M n=7, mC-Salm F n=15, mI-Salm M n=12, mI-Salm F n=18). (E) Frequency and numbers of Treg cells, Tr17 cells and Th17 cells in spleen. (mC-Salm M n=13, mC-Salm F n=17, mI-Salm M n=7, mI-Salm F n=14). Data in **B**-**E** are from 2-4 independent experiments using 3-4 pregnant dams per group and 4-6 offspring per pregnancy. One-way ANOVA adjusted P: ‘ns’ not significant, * <0.033, ** < 0.0021, *** < 0.0002, **** <0.0001. Summary of statistical analyses are shown in **Supplemental Data 1**.

### Increased MMCs in male offspring of previously infected dams correlates with increased splenic RORγt^+^ T cells

Maternal infection during pregnancy did not alter the distribution of major immune subsets in the spleen of offspring (**Supp Fig. 4**). An intestinal tissue-specific T helper 17 effector response to postnatal microbial exposure was previously reported in offspring of prenatally infected dams(26). However, we did not observe a significant increase in RORγt expressing CD4^+^ T (Th17) cells in the small intestine of 4-5 weeks old offspring (**Supp Fig. 5A**). The genetic background of animals influences gut microbial diversity(31) that likely has an impact on Th17 cell homeostasis. The mixed B6-Balb/c background of offspring of allogeneic matings as well as the earlier timing of maternal infection at E7.5 in our model may contribute to the lack of a gut specific Th17 response as seen previously. However, maternal infection was indeed associated with a significant increase in RORγt^+^ and IL-17^+^ Th17 cells in the spleen of male offspring compared to those from uninfected dams (**Fig. 2D, Supp Fig. 5B-D**). There were no differences in IFNg and IL-4 producing Th1 and Th2 cells respectively (**Supp Fig. 5B-D**). There was also a significant positive correlation between MMCs and Th17 cells in the spleen of male, but not female offspring (**Fig. 2E, Supp Fig. 5E**). While distribution of CD4^+^FOXP3^+^ Treg cells was unaffected, the frequency of RORγt expressing Treg cells (Tr17), that are induced by microbes in the gut as well as by immunization in peripheral lymph nodes(32–34), was also increased and correlated with frequency of MMCs (**Fig. 2D, E**). There were no changes in other RORγt expressing cell subsets including TCRb^neg^CD3^neg^RORγt^+^ cells, iNKT17 cells and CD11c^+^RORγt^+^ dendritic cells (**Supp Fig. 5B, F**). Thus, transient pregnancy-associated maternal infection influenced the homeostasis of RORγt expressing CD4^+^ T cells in offspring that correlated with the frequency of MMCs.

**Figure 4:**
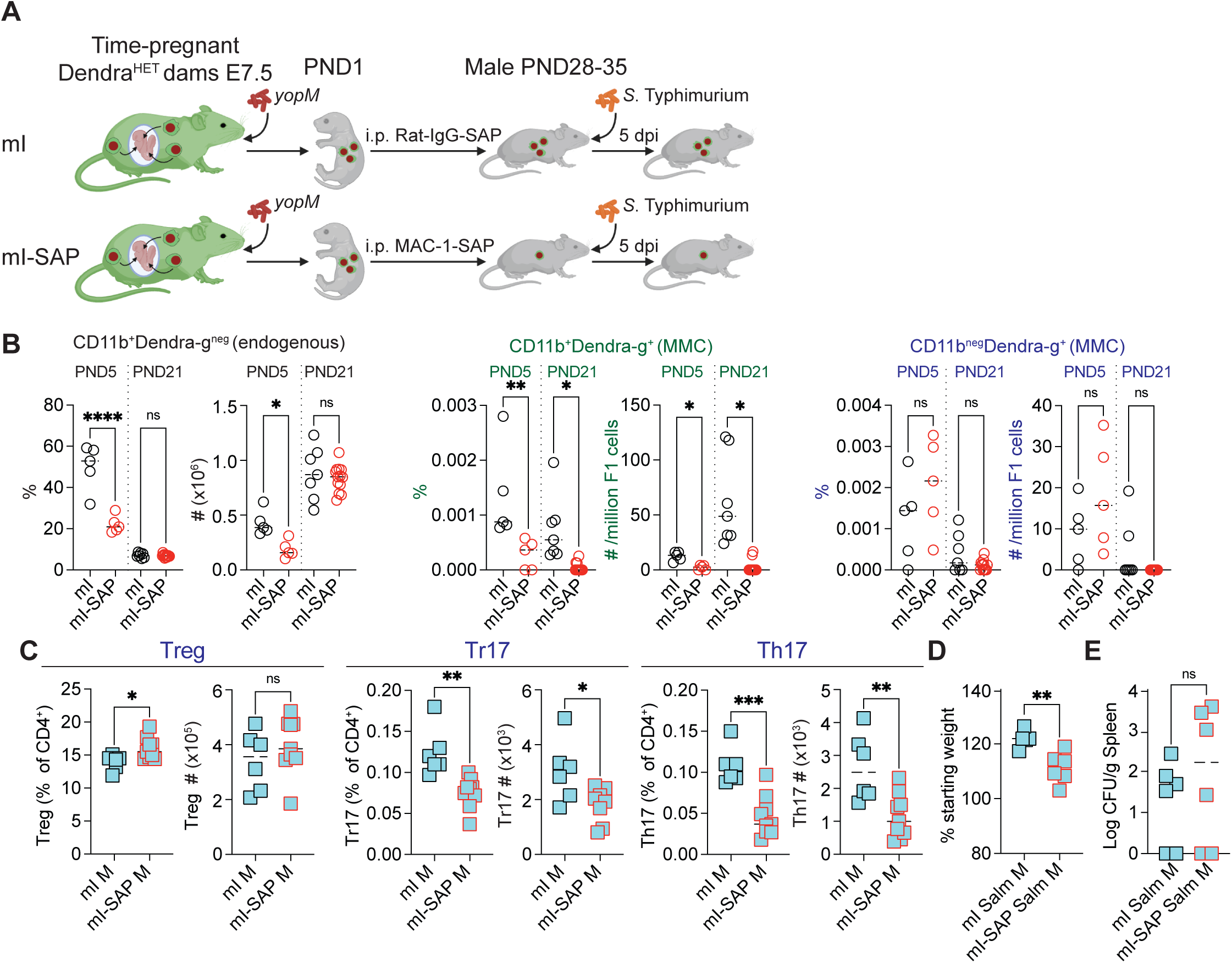
Loss of CD11b^+^ MMCs alters type 17 T effector cell profile and susceptibility to *Salmonella* infection in male offspring. **(A)** Newborn (PND0) Dendra^WT^ offspring of previously infected dams (mI) were injected with control Rat-IgG SAP (mI Ctrl) or MAC1-SAP (mI SAP) antibody to deplete CD11b^+^ cells. **(B)** Frequency and numbers of (*Left*) endogenous (offspring origin) Dendra^neg^CD11b^+^ cells, (*Middle*) CD11b^+^ MMCs and (*Right*) CD11b^neg^ MMCs in PND5 and PND21 offspring of previously infected dams that were treated with either control rat-IgG-SAP (mI) or Mac-1-SAP (mI-SAP) antibody. (PND5: mI n=5, mI-SAP n=5; PND21: mI n=7, mI-SAP n=13). **(C)** Frequency and numbers of splenic Treg cells, Tr17 cells and Th17 cells in control and Mac- 1-SAP treated male PND28-35 offspring of previously infected dams. (mI M n=6, mI-SAP M n=9). Data in **B, C** are from 2-4 independent experiments using offspring of 3-4 infected dams in each group. One-way ANOVA adjusted P: ‘ns’ not significant, * <0.033, ** < 0.0021, *** < 0.0002, **** <0.0001. **(D)** Percent change in body weight 5 days after *Salmonella* infection in male offspring. (mI Salm M n=6, mI-SAP Salm M n=6). **(E)** Splenic bacterial burden 5 days after *Salmonella* infection. (mI Salm M n=6, mI-SAP Salm M n=6). Data in **D** and **E** are from 2 independent experiments using offspring of 3-4 infected dams in each group. t-test P: ‘ns’ not significant, * <0.033, ** < 0.0021, *** < 0.0002, **** <0.0001. Summary of statistical analyses are shown in **Supplemental Data 1**.

Maternal IL-6 produced during pregnancy-associated infection can induce intestinal Th17 responses in offspring(26). To exclude the contribution of trans-placentally transferred soluble factors including cytokines and antibodies, and to specifically determine if maternal cells regulated Th17 cell homeostasis in offspring lymphoid tissues, we conducted an adoptive transfer experiment (**Fig. 2F**). We focused on CD11b^+^ MMCs as their frequency was increased in male offspring of infected dams at the fetal and immediate postnatal stages (**Fig. 1**). Sorted Dendra-g^+^ CD11b^+^ and CD11b^neg^ splenocytes from Dendra^HET^ pregnant females 5 days after *yopM* infection were intravenously injected into timed pregnant C57BL/6 females at E7.5. Dendra-g^+^CD11b^+^ cells sorted from uninfected pregnant females were used as controls. There were no differences in the frequency of transferred maternal cells acquired by offspring across groups (**Fig. 2G**). However, offspring of B6 females that received CD11b^+^ cells from previously infected mothers showed elevated Tr17 and Th17 cells in the spleen compared to offspring of recipients of CD11b^+^ cells from control uninfected mothers (**Fig. 2H**). The ability to induce RORγt^+^ T cell responses was restricted to the CD11b^+^ population as offspring of recipients of CD11b^neg^ cells from previously infected mothers were unable to do so. Further, the effect of acquired CD11b^+^ cells was specific to RORγt^+^ T cells as the frequency of Treg cells remained unchanged. These data suggest that transient maternal infection during pregnancy enhanced the ability of MMCs to promote a type 17 T effector cell response in the offspring.

### Maternal infection increases resistance of male offspring to *Salmonella* Typhimurium infection

We next sought to determine if there were functional consequences of the association between MMCs from previously infected mothers and the male offspring’s own Th17 response. IL-17 producing RORγt^+^ cells have been shown to be protective in *Salmonella* infection(35). We, therefore, challenged male and female offspring of previously infected and control dams with *Salmonella* Typhimurium at postnatal day 25-30 (PND25-30) (**Fig. 3A**). Male offspring of previously infected dams had significantly improved survival following acute *Salmonella* infection than the male offspring of uninfected dams (**Fig. 3B**). They lost significantly less weight and displayed a small but statistically significant decrease in splenic pathogen burdens (**Fig. 3C, D, Supp.** Fig. 6). These differences in handling *Salmonella* were associated with significantly higher Tr17 and Th17 cells, but not Treg cells, in the male offspring of infected mothers (**Fig. 3E**).

### Depletion of CD11b^+^ MMCs alters Th17 cell homeostasis and susceptibility to infection in male offspring

Data so far pointed to a role for maternally conditioned CD11b^+^ MMCs in regulating Type 17 T cell responses in male offspring. To determine whether CD11b^+^ MMCs are required for this phenomenon, we depleted MMCs in newborn male mice using a saporin-conjugated antibody to Mac-1 (CD11b)(36). Male offspring of previously infected dams were administered either Mac-1- SAP or rat-IgG-SAP antibody on the day of birth (**Fig. 4A**). Mac-1-SAP treatment is expected to kill all CD11b^+^ cells in newborn mice, including CD11b expressing maternal cells and the offspring’s own CD11b expressing myeloid cells. However, we reasoned that while gestationally transferred MMCs would be irreversibly killed, the offspring’s own CD11b^+^ compartment would be replenished from new precursors arising in the bone marrow. Endogenous CD11b^+^ cells were indeed significantly reduced 5 days after Mac-1-SAP treatment, but their numbers recovered by 3 weeks (**Fig. 4B**). CD11b^+^ MMCs were also reduced at 5 days post treatment, but unlike endogenous CD11b^+^ cells, remained significantly reduced at 3 weeks (**Fig. 4B**). There were no differences in CD11b^neg^ MMCs at either timepoint (**Fig. 4B**). The decrease in CD11b^+^ MMCs was accompanied by a significant decrease in Tr17 and Th17 cells at PND21 in Mac1-SAP treated male offspring of previously infected dams (**Fig. 4C**). Finally, we challenged Mac-1-SAP and control rat-IgG-SAP treated offspring with *Salmonella* Typhimurium at postnatal day 25-30 (PND25-30). Despite splenic bacterial burden not being significantly different at day 5 after infection, Mac-1-SAP treated male offspring of previously infected dams appeared to do more poorly and exhibited greater weight loss compared to control rat-IgG-SAP treated male offspring after *Salmonella* infection (**Fig. 4D, E**).

## Discussion

Vertically transferred maternal factors play crucial roles in regulating offspring immunity in early life. Here, we showed that pregnancy-associated transient infection influenced MMC homeostasis and function in offspring. Increased MMCs specifically in male offspring of previously infected mothers promoted a RORγt expressing effector T cell (type 17) response that decreased susceptibility to infection. These findings highlight a sex-differential function of MMCs that is revealed upon maternal infection during pregnancy.

Allogeneic MMCs induce Treg cells in female offspring to establish tolerance to non-inherited maternal antigens (NIMA) and enable their persistence into adulthood. In our studies, we consistently observed increased frequency of MMCs in young-adult female offspring of naïve mothers compared to male offspring, suggesting distinct sex-specific mechanisms that maintain MMCs in the long term. Maternal infection during pregnancy, however, altered this dynamic and led to an induction of MMCs in male offspring of infected mothers compared to those from naïve mothers. Maternal cells in male offspring induced a pro-inflammatory Tr17/Th17 program in lymphoid tissues that conferred protection from severe *Salmonella* infection in young adulthood. The mechanisms by which MMCs induce Type 17 responses is yet unknown. Given the phenotypic resemblance of MMCs to CD11b^+^Ly6G^+^Ly6C^+^ MDSCs, possible mechanisms of action could involve production of arginase-1 and reactive oxygen species that have been shown to regulate Th17 differentiation in models of systemic lupus erythematosus and lupus nephritis (37, 38). Of note, there was a significantly greater impact on overall health and body weight than on control of bacterial load in infected offspring. Disease tolerance is a phenomenon whereby the host prioritizes tissue damage control over elimination of pathogen as a means to preserve long-term homeostasis(39). It is intriguing to speculate that MMCs safeguard their own survival and the reproductive fitness of their female host by inducing NIMA-specific Treg cells but may influence the state of disease tolerance and infection risk in their male hosts by inducing a type 17 T cell response. Sex differences in immunity lead to enhanced vaccine responses, greater resistance to infection and an increased risk of autoimmune conditions in female relative to males(40). Newborn males are particularly vulnerable to infections and death compared to females(41). However, increased MMCs in male offspring correlates with reduced early life infections in humans(18).

Our findings extend this observation to include a role for the maternal immunological experience that influences acquired MMCs to provide immune benefits to male offspring in infectious settings.

Sex-specific differences in tissue immunity have been reported. For example, type 2 immunity is lower in male mice compared to female mice after allergen exposure in a model of allergic airway inflammation(42). It is noteworthy that we observed an impact of MMCs on male offspring T effector phenotype specifically in the spleen but not at tissue sites such as the small intestine. While MMCs have been reported in multiple tissues(4), they appear to be enriched at sites of hematopoiesis including the fetal bone marrow(18) and, in our studies, the yolk sac and fetal liver. Further, in contrast to their presence in the female uterus, they are notably absent in male reproductive tissues of the prostrate(17). MMCs may thus selectively take up residence in specific tissues where they serve as an alloantigen source to influence T cell immunity. We also observed that sex differences in MMC numbers only manifested during the postnatal period and were absent during fetal development. One possibility is that fluctuations in sex-specific hormone levels influenced by age and external environmental factors such as the microbiota may differentially regulate the homeostasis of maternal cells in male and female offspring(43, 44). Androgens and androgen receptor signaling were recently shown to regulate skin T cell expansion in a sex specific manner(45). Future studies should uncover the role of sex hormones in regulating MMC homeostasis.

Infection induced maternal immune activation is associated with increased expression and trans- placental transport of inflammatory cytokines that influence immune functionality and offspring disease susceptibility. Vertically transferred maternal IL-6 is one conveyor of maternal inflammation to the developing fetus, affecting functional brain networks in the immediate postnatal period(46). Maternal IL-6 also imprints altered functionality into the developing fetal intestine that influences immune responses at barrier surfaces in adulthood(26) However, in our studies, we showed that the transfer of CD11b^+^ cells from infected mothers into naïve pregnant recipients and their subsequent acquisition by offspring was sufficient to induce the type 17 program in offspring lymphoid tissue. A direct role for trans-placental cytokines to influence this T cell phenotype is further negated by the developmental timing of T cell appearance in murine offspring. The mouse fetal thymus is notably immature and unlike its human counterpart, is unable to support T cell development until 4-5 days after birth(47). Thus, inflammatory maternal cytokines during gestation are unlikely to directly program an effector T cell phenotype that arises later in ontogeny. Future studies will need to address whether maternal infection alters the acquisition and/or persistence of MMCs in offspring and how it imprints distinct functionality in subsets of MMCs, including the CD11b^+^ MMCs, that induces the type 17 T cell effector program in lymphoid tissues of male offspring.

### Limitations of study

In proof-of-concept studies, we tracked MMCs in animal models and uncovered their sex differential functions in offspring following transient maternal infection with *yopM*. Maternal immune activation following infection during pregnancy can take different forms depending on the type and severity of infection as well as the involvement of the placenta that may differentially influence acquired maternal cells and their function. Studies using additional challenges combined with consideration of timing of maternal infection as well as maternal parity are warranted to clarify their impact on MMCs. All mice used in our experiments were on the B6 or B6 x Balb/c background and so carried the Nramp1 nonfunctional allele that confers susceptibility to *Salmonella* infection. B6 mice are widely used in studies of salmonellosis and have provided several important insights into pathogenesis and the anti-*Salmonella* immune response. Nevertheless, we acknowledge that our findings must be interpreted with the understanding that they may have been influenced by Nramp1 functionality.

Our study focused on trans-placentally transferred maternal cells and the impact of their depletion on offspring susceptibility to infectious challenge. However, MMCs are also transferred through breastmilk in early life, and we might have missed defining a role for these cells, particularly in regulating mucosal immunity in offspring. MMCs are a rare population of cells and their study in humans have been limited by available methods to identify them and establish causal relationships in disease settings. We acknowledge that developmental timelines are distinct in humans and mice and while it may be premature to draw direct parallels between our findings and observations made in humans, our studies clearly call for dedicated exploration of maternal immune challenges and their impact on MMCs.

## Acknowledgments

We are thankful for the resources of the Center for Computational and Integrative Biology and the Mucosal Immunology and Biology Research Center at MGH that supported this study. NJ was supported by grants R21AI168518, R01AI154626.

## Author contributions

BJ, DK SK, HM, ES performed experiments and analyzed data with help from LL, PK, CP. NJ, BC and BS designed experiments and NJ wrote the paper with input from all authors.

## Declaration of Interests

The authors declare no competing interests.

**STAR Methods**

## RESOURCE AVAILABILITY

### Lead Contact

Further information and requests for resources and reagents should be directed to and will be fulfilled by the lead contact, Nitya Jain.

### Materials availability

This study did not generate new unique reagents.

### Data and code availability

All data reported in this paper will be shared by the lead contact upon request. This paper does not report original code. Any additional information required to reanalyze the data reported in this paper is available from the lead contact upon request.

## EXPERIMENTAL MODEL AND STUDY PARTICIPANT DETAILS

### Mice

Animal studies were conducted under protocols approved by the Institutional Animal Care and Use Committee of Massachusetts General Hospital. The following animal lines were used: PhAM^excised^ mice (B6;129S-Gt(ROSA)26Sor^tm1.1(CAG-COX8A/Dendra2)Dcc^/J; Jax strain 018397), C57BL/6 mice (Jax strain: 00664) and BALB/cJ mice (Jax strain 000651). PhAM^excised^ (Dendra^HOM^) mice were mated with C57BL/6 mice to generate Dendra^HET^ females that were used as breeders in allogeneic matings with Balb/cJ mice. Mice were housed in HPPF facilities (*Heliobacter-* and *Pasteurella pneumotropica-* free) before mating and transferred to BL-2

facilities for infection. Sex and age of all mice used are reported in figure legends. *Yersinia pseudotuberculosis yopM* strain was a gift from Dr. Igor Brodsky and Dr. James Bliska(27). The SL1344 strain of *Salmonella* Typhimurium were a gift from Dr. Beth McCormick.

## METHOD DETAILS

### Timed matings

Littermate female Dendra^HET^ mice from a Dendra^HOM^ and C57BL/6 cross were used in all experiments. Singly housed experienced Balb/cJ male mice were mated with Dendra^HET^ females that were in proestrus or estrus. Presence of vaginal plug was monitored over the next 3-4 days and males were separated from the breeding cage once a plug was observed. The morning on which plugs were observed was designated as embryonic day 0.5. Offspring were weaned at 3 weeks after birth and age-matched male and female mice were used in experiments as specified.

### Yersinia pseudotuberculosis infection

The *yopM* mutant strain of *Y. pseudotuberculosis* 32777 was grown in 2 mL of YT Broth (2X) (Sigma) at 30°C with shaking at 250 rpm. 100 µL of overnight culture was diluted tenfold with phosphate-buffered saline (PBS) and the OD value obtained with a spectrophotometer absorbance reading at 600nm. Overnight cultures were then appropriately diluted with PBS to reach an inoculum size of ∼ 2 x 10^7^ colony forming units (CFUs) / 100 µL. Time-pregnant female mice were administered 100 µL of bacterial suspension by oral gavage. Stool samples collected from infected pregnant dams were weighed, and fecal DNA extracted using the QIAmp Fast DNA Stool Mini Kit (Qiagen). The *yscF* gene was quantified against a standard curve by quantitative PCR (forward primer 5′-ATGAGTAACTTCTCTG- GATTTACG-3′, reverse primer 5′-TTATGG-GAACTTCTGTAGGATG-3′). Quantitative PCR was performed using the QuantStudio 3 Real- Tine PCR system with a well-volume of 20 µL.

### *Salmonella* Typhimurium infection

The SL1344 strain of *S.* Typhimurium was cultured in LB with 50 µg/mL streptomycin at 37°C while shaking at 250 rpm. After 8 hours, 10 µL of the shaking culture was back diluted into 10 mL of fresh LB with 50 µg/mL streptomycin and incubated overnight without shaking. Overnight standing cultures were centrifuged and diluted to 2 x 10^7^ CFU per 100 µL in PBS for infection. 28-35 days old offspring of Dendra^HET^ x Balb/cJ matings were infected by oral gavage. At 5 days after infection, harvested tissues were weighed and homogenized in 0.5% Triton in PBS. Serial dilutions were spread plated on LB agar, incubated overnight at 37°C before calculating CFUs.

### Tissue collection and single cell preparation

Single cell suspensions were prepared from the spleens of newborn and adult animals following mechanical homogenization as described previously(48). RBCs were lysed using ACK lysis buffer. Cells from the small intestine lamina propria were isolated after removing epithelial fraction followed by digestion in Liberase (Roche). Embryonic tissues (FL and YS) were homogenized between frosted glass slides and digested in Liberase TL (thermolysin low) as described before(49). Cells isolated from all tissues were counted on a Countess 3 (Thermo Fisher Scientific) instrument and resuspended in FACS buffer in preparation for staining.

### *In vitro* cell stimulation

Splenocytes were cultured with 1X protein transport inhibitor cocktail (Thermo Fisher Scientific)

and 1X cell stimulation cocktail (Thermo Fisher Scientific) for 16 hours. Cultures were harvested and cells stained for surface markers. Cells were then fixed and permeabilized with BD Cytofix/Cytoperm buffer and stained with cytokine antibodies before analysis on a flow cytometry.

### Flow Cytometry

Antibody and reagent information is provided in **Table S1**. A maximum of 3 x 10^6^ cells were stained with fluorophore-conjugated antibodies. After surface staining, cells were fixed and permeabilized using the Foxp3/Transcription Factor Staining Buffer Set (Thermofisher Scientific) and stained with intracellular transcription factor antibodies. Dead cells were excluded using LIVE/DEAD Fixable Blue Dead Cell Stain Kit (Invitrogen Life Technologies). Samples were acquired on a 5 laser Cytek Aurora spectral analyzer. Single color controls were made using UltraComp eBeads for spectral unmixing on the cytometer.

Flow cytometry-based genotyping was performed to identify Dendra^WT^ and Dendra^HET^ samples for embryonic and newborn timepoints. Single cell suspensions from the fetal liver were acquired on the Cytek in plate mode and Dendra^HET^ samples, in which >98% of total events are Dendra-g^+^, were identified and discarded. Only stained Dendra^WT^ samples were acquired and the ‘sample recovery’ function and a 30 second water wash between each sample was programmed into the SpectroFlo software that minimized carryover(50). Single cell suspensions from entire organs at the fetal and postnatal day 1 time points were run through the cytometer. For adult tissues, up to 1 x 10^6^ cells were acquired at a flow rate between 50 to 70 uL / minute. The data were analyzed using FlowJo v10 software (BD).

### *In vivo* manipulations

2-5 x10^4^ MACS enriched CD11b^+^ and CD11b^neg^ cells (Miltenyi Biotec) from maternal spleens were i.v. injected into recipient females. Single dose of Mac-1-SAP or Rat-IgG-SAP (ATS Bio; 4 micrograms per animal) was i.p. injected into newborn mice.

### Statistical Analysis

GraphPad Prism (v.10.4.1) were used to perform data analyses. Data from all statistical analyses performed are shown in **Supplemental Data 1**. Two-tailed unpaired *t* tests were used for comparisons between groups at similar timepoints. Comparisons between male and female offspring of infected and control dams were performed using One-way ANOVA with Tukey’s multiple comparisons test with a single pooled variance. All sex differences were established using Two-way ANOVAs combined with Tukey’s multiple comparison test. No randomization was performed, and investigators were not blinded to group allocations.

### Supplemental Data 1

Summary of statistical analyses used in manuscript, related to Figures 1-4 and Supp Figs. 1-6.

---

**Supplemental Figure 1:**
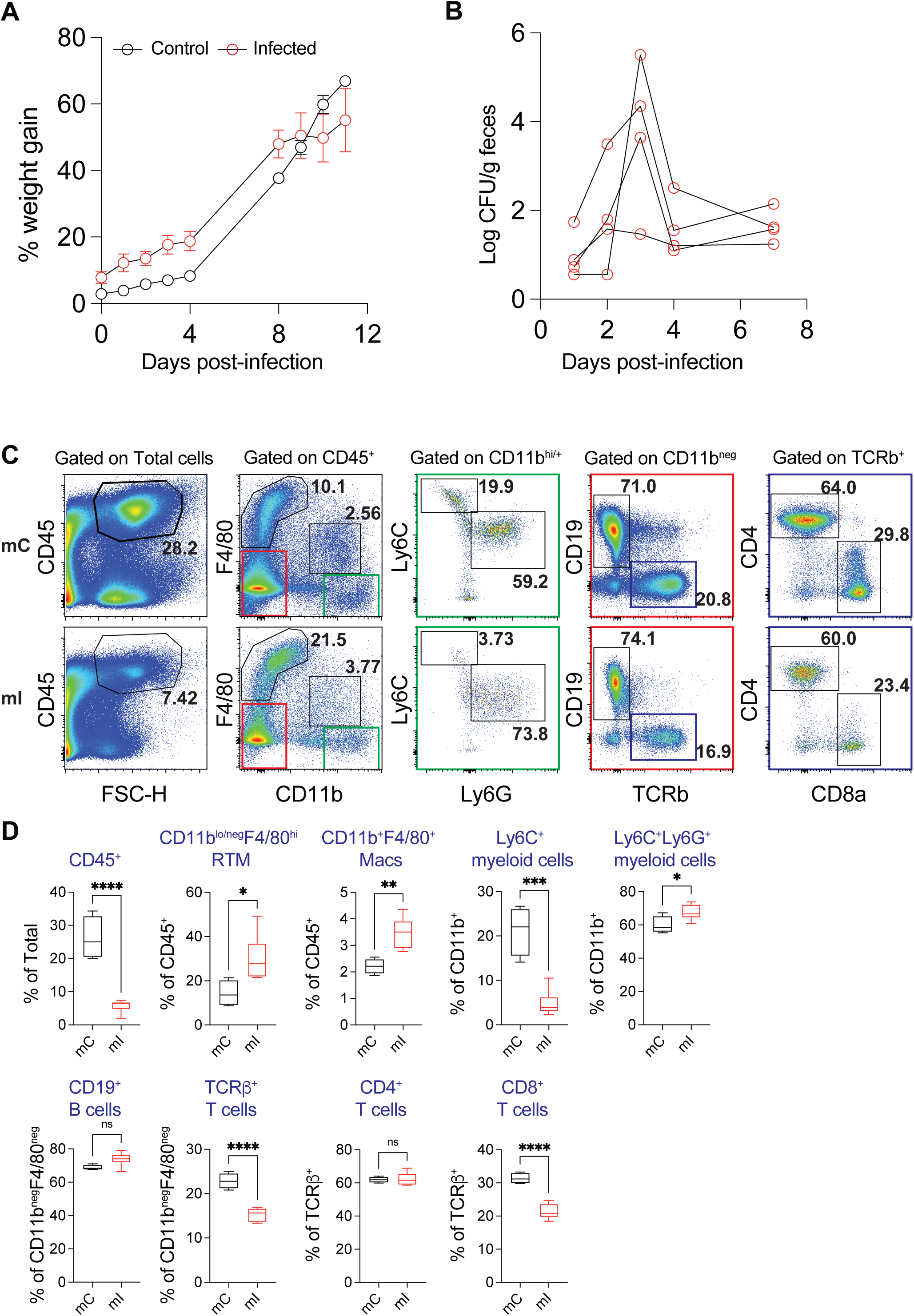
*yopM* models of maternally restricted transient infection, related to Figure 1 Timed-pregnant Dendra^HET^ dams were orally administered *Y. pseudotuberculosis yopM* at embryonic day 7.5 (E7.5). (A) Body weight of control (mC) and *yopM*-infected (mI) dams over the course of pregnancy. (mC n=3; mI n=5) (B) yopM burden in feces at 1, 2, 3, 4 and 7 days after infection (n=5). (C) Maternal splenic immune cell distribution 5 days after *yopM* infection. Representative flow cytometry dot plots show gating strategy to identify immune subsets in mC and mI females. (D) Frequency of CD45^+^ cells, resident tissue macrophages (RTMs; CD11b^lo/neg^F4/80^hi^), macrophages (Macs; CD11b^+^F4/80^+^), CD11b^+^Ly6C^+^ myeloid cells, CD11b^+^Ly6C^+^Ly6G^+^ myeloid cells, CD19^+^ B cells, TCRb^+^ T cells, CD4^+^T cells and CD8^+^ T cells. (mC n=4, mI n=6; data are representative of 2 independent experiments). Summary of statistical analyses are shown in **Supplemental Data 1**.

**Supplemental Figure 2:**
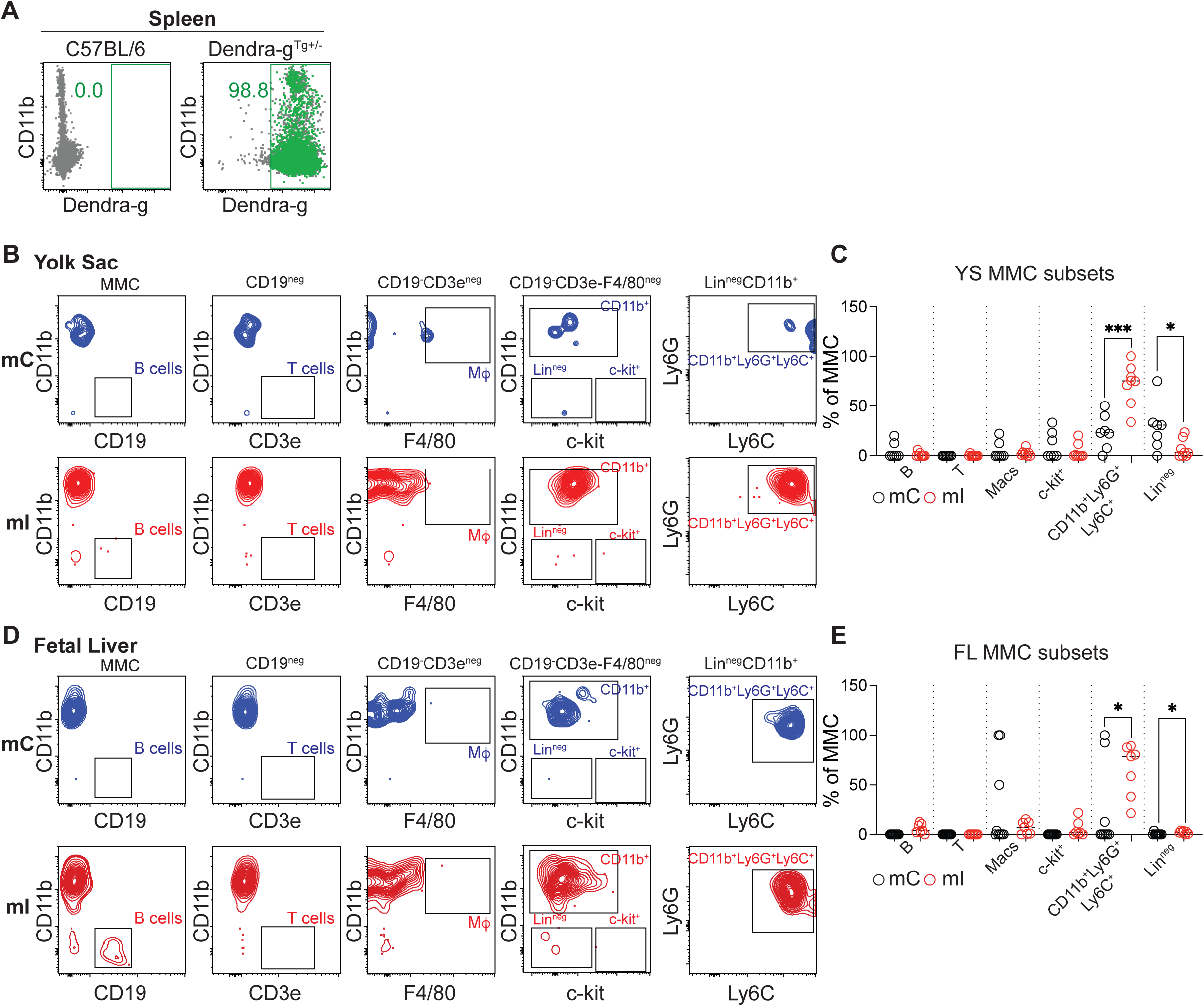
Gating strategy to identify MMC subsets in E14.5 YS and FL, related to Figure 1. (A) Splenocytes from C57BL/6 and Dendra-g^Tg+/-^ were analyzed by flow cytometry to set Dendra- g^+^ gate. **(B-D)** Allogeneic matings of Dendra^HET^ females with BALB/cJ males were set up to generate Dendra^WT^ offspring. Timed-pregnant Dendra^HET^ dams were orally administered *Y. pseudotuberculosis yopM* at embryonic day 7.5 (E7.5). Yolk sac (YS) and fetal liver (FL) tissues from E14.5 fetuses from infected (mI) and control (mC) mothers were analyzed by flow cytometry. (B) **Identification of MMCs in YS, FL:** (*Left*) Representative dot plots show expression of Dendra-g (x-axis) and CD11b (y-axis) on CD45^+^ cells; green: Dendra-g^+^ cells, gray: Dendra-g^neg^ cells. (*Right*) Dendra-g^+^ cells were further analyzed for expression of MHC haplotypes H2-Kd (Balb/c) and H2-Kb (B6). MMCs of maternal B6 origin were identified as H2-Kd^neg^H2-Kb^+^. (C) **Subsets of MMCs in YS**: Dendra-g^+^ cells of maternal B6 origin (H2-Kd^neg^H2-Kb^+^) in YS were analyzed for expression of lineage markers to identify CD19^+^ B cells, CD3e^+^ T cells, CD11b^+^F4/80^+^ macrophages and c-kit^+^ cells. CD11b^+^F4/80^neg^ cells were analyzed further for expression of Ly6G and Ly6C to identify CD11b^+^Ly6C^+^Ly6G^neg^ cells. (D) Frequency of MMC subsets in mC and mI E14.5 YS (mC: n=7, mI: n=8). Error bars are SEM. (E) **Subsets of MMCs in FL:** Dendra-g^+^ cells of maternal B6 origin (H2-Kd^neg^H2-Kb^+^) in FL were analyzed as in (**C**). (F) Frequency of MMC subsets in mC and mI E14.5 FL (mC: n=9, mI: n=7). Error bars are SEM. Data in B-F are representative of 2 independent experiments.

**Supplemental Figure 3:**
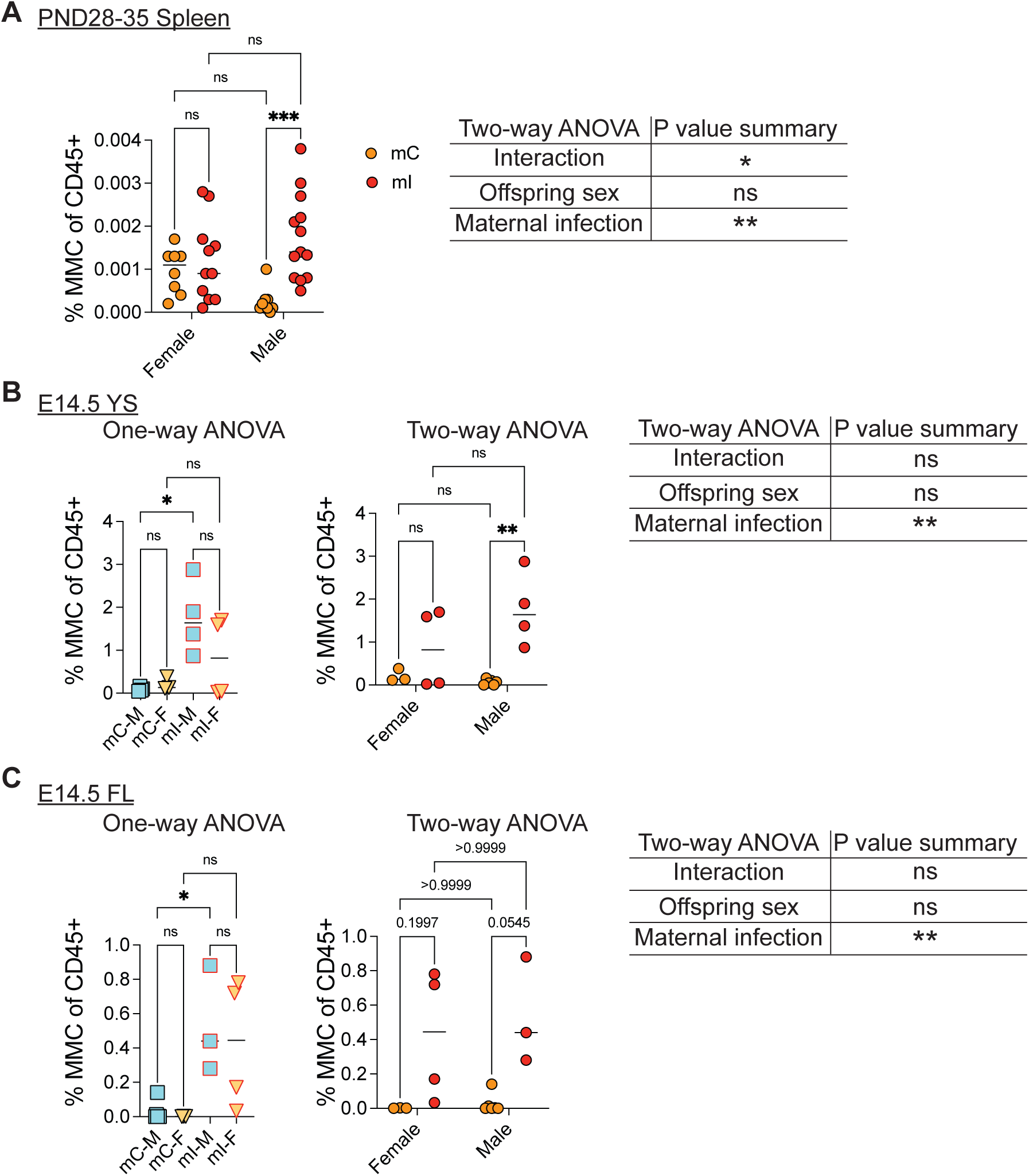
Pregnancy-associated maternal infection increases acquisition of MMCs in male fetuses, related to Figure 1 Allogeneic matings of Dendra^HET^ females with BALB/cJ males were set up to generate Dendra^WT^ offspring. Timed-pregnant Dendra^HET^ dams were orally administered *Y. pseudotuberculosis yopM* at embryonic day 7.5 (E7.5). Male and female Dendra^WT^ offspring of infected (mI) and control (mC) mothers were analyzed at PND28-35 and E14.5. (A) Frequency of MMCs within CD45^+^ cells in offspring spleens of previously infected dams segregated by sex. Two-way ANOVA results are shown on the right. (mC male n=9, mC female n=8, mI male n=13, mI female n=9; data are from 5 independent experiments using 2-3 pregnant mice dams per group and 2-5 offspring per pregnancy). (B) E14.5 Yolk Sac (YS): (*Left*) Frequency of MMCs within CD45^+^ cells. One-way ANOVA analysis is shown. (*Right*) Frequency of MMCs within CD45^+^ cells in offspring spleens of previously infected dams segregated by sex. Two-way ANOVA analysis is shown. (mC M n=4, mC F n=3, mI M n=4, mI F n=4; data are from 2 independent experiments using 2 pregnant mice dams per group and 3-5 offspring per pregnancy). (C) E14.5 Fetal Liver (FL): (*Left*) Gating on CD45^+^ cells identifying Dendra-g^+^ MMCs in male and female fetuses of mC and mI dams. (*Middle*) Frequency of MMCs within CD45^+^ cells. One- way ANOVA analysis is shown. (*Right*) Frequency of MMCs within CD45^+^ cells in offspring spleens of previously infected dams segregated by sex. Two-way ANOVA analysis is shown. (mC M n=6, mC F n=3, mI M n=3, mI F n=4; data are from 3 independent experiments using 3-5 pregnant mice dams per group and 3-5 offspring per pregnancy.) ANOVA adjusted P * <0.033, ** < 0.0021, *** < 0.0002, **** <0.0001 Summary of statistical analyses are shown in **Supplemental Data 1**.

**Supplemental Figure 4:**
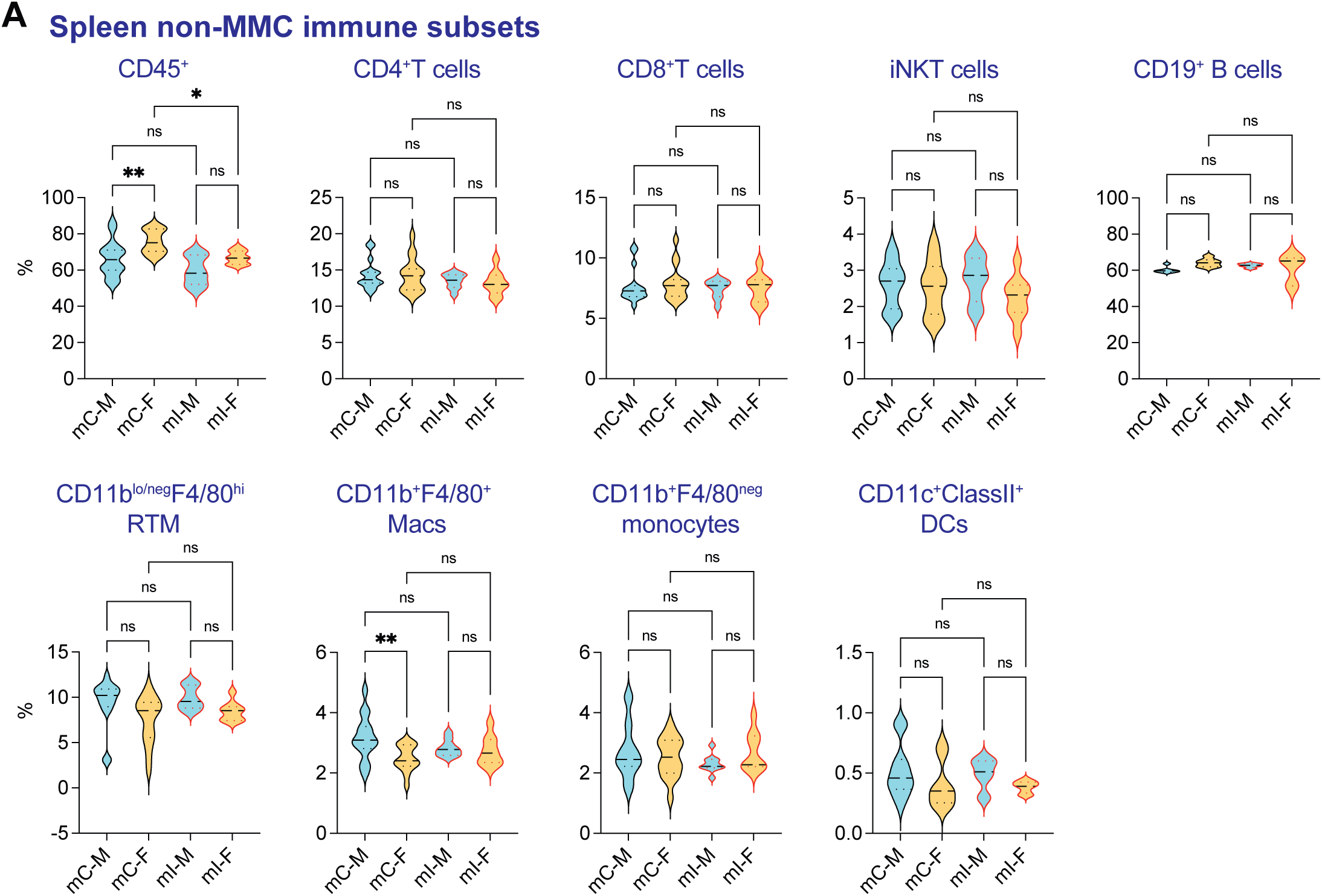
Distribution of splenic [non-MMC] immune subsets in spleen of PND28-35 offspring, related to Figure 2 Allogeneic matings of Dendra^HET^ females with BALB/cJ males were set up to generate Dendra^WT^ offspring. Timed-pregnant Dendra^HET^ dams were orally administered *Y. pseudotuberculosis yopM* at embryonic day 7.5 (E7.5). Male and female Dendra^WT^ offspring of infected (mI) and control (mC) mothers were analyzed at PND28-35. **(A)** Frequency of CD45^+^ cell, CD4^+^T cells, CD8^+^ T cells, iNKT cells, CD19^+^ B cells, resident tissue macrophages (RTMs; CD11b^lo/neg^F4/80^hi^), macrophages (Macs; CD11b^+^F4/80^+^), CD11b^+^F4/80^neg^ monocytes and CD11c^+^ClassII^+^ dendritic cells. (mC male n=16, mC female n=16, mI male n=9, mI female n=9; data are from 5 independent experiments using 3-4 pregnant mice dams per group and 4-6 offspring per pregnancy) One-way ANOVA adjusted P: ns not significant, * <0.033, ** < 0.0021, *** < 0.0002, **** <0.0001. Summary of statistical analyses are shown in **Supplemental Data 1**.

**Supplemental Figure 5:**
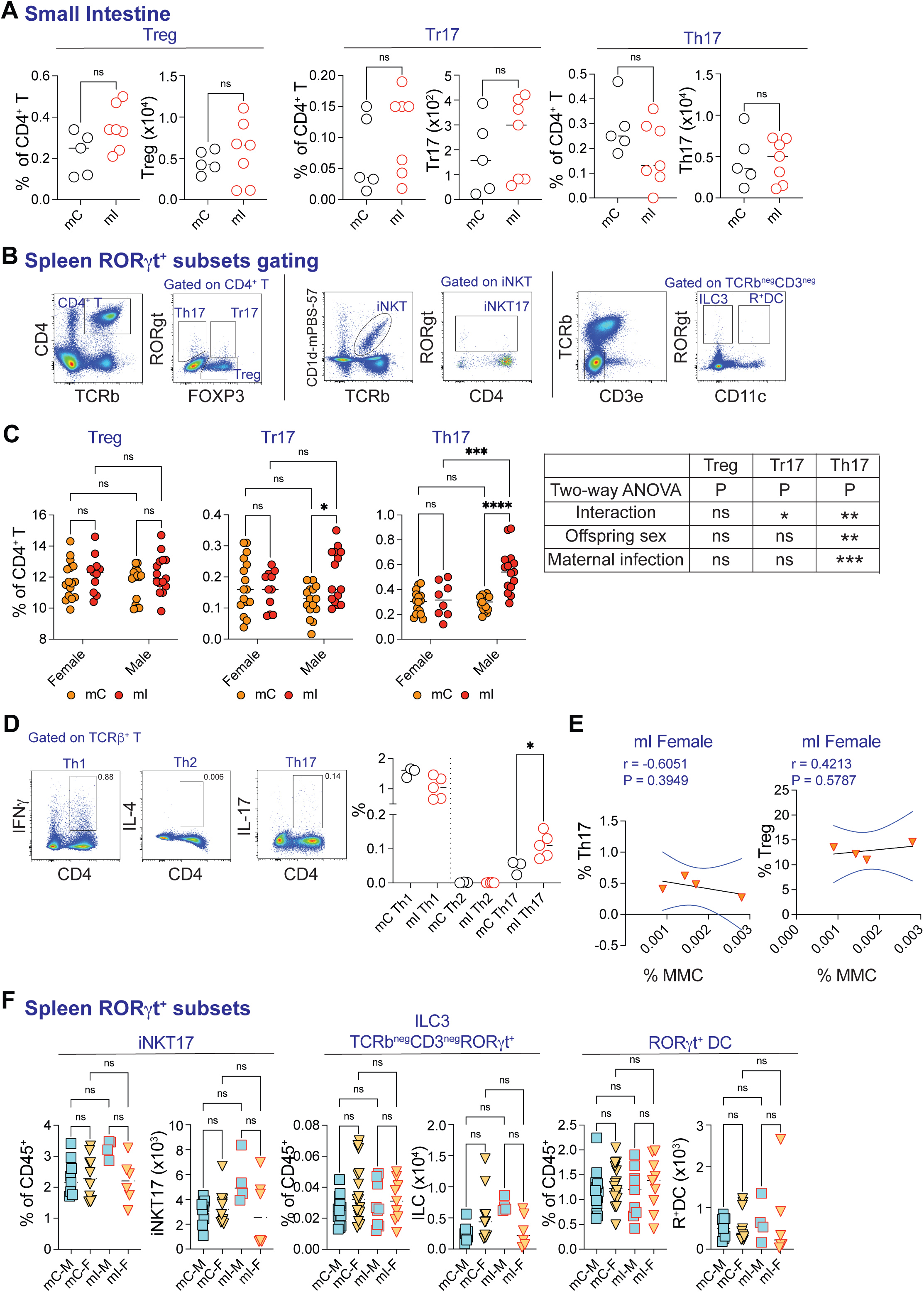
Distribution of RORγt expressing cell subsets in small intestine and spleen of PND28-35 offspring, related to Figure 2 Allogeneic matings of Dendra^HET^ females with BALB/cJ males were set up to generate Dendra^WT^ offspring. Timed-pregnant Dendra^HET^ dams were orally administered *Y. pseudotuberculosis yopM* at embryonic day 7.5 (E7.5). Male and female Dendra^WT^ offspring of infected (mI) and control (mC) mothers were analyzed at PND28-35. (A) **Small Intestine (SI):** Frequency and numbers of (*From left to right*) Treg cells, RORγt^+^ Treg (Tr17) cells and Th17 cells. (mC n=5, mI n=7; data are from 2 independent experiments using 2 pregnant dam per group and 2-4 offspring per pregnancy; t-test P ‘ns’: not significant). (B) **Spleen RORγt^+^ subsets gating strategy:** Representative flow cytometry dot plots show gating strategy to identify RORγt^+^ subsets including CD4^+^RORγt^+^ Th17, CD4^+^RORγt^+^FOXP3^+^ Tr17, CD4^+^RORγt^neg^FOXP3^+^ Treg, CD1d-mPBS57-tetramer^+^TCRb^+^RORγt^+^ iNKT17, CD3e^neg^TCRý^neg^RORγt^+^ ILC3 and CD11c^+^RORγt^+^ DCs. (C) **Spleen PND28-35:** Frequency of Tregs, Tr17 and Th17 cells in offspring spleens of previously infected dams segregated by sex. Two-way ANOVA analysis is shown (mC male n=15, mC female n=16, mI male n=15, mI female n=11; data are from 5 independent experiments using 3-4 pregnant mice dams per group and 4-6 offspring per pregnancy; two-way ANOVA adjusted P: ns not significant, * <0.033, ** < 0.0021, *** < 0.0002, **** <0.0001). (D) (*Left*) Gating strategy for identifying CD4^+^IFNg^+^ Th1 cells CD4^+^IL-4^+^ Th2 cells and CD4^+^ IL-17^+^ Th17 cells. Splenocytes from C57BL/6 mice were stimulated with PMA + Ionomycin in the presence of Golgi Stop and Golgi Plug for 6 hours followed by intracellular cytokine staining. (*Right*) Frequency of Th1, Th2 and Th17 cells in splenocytes from mC and mI offspring (6-8 weeks old) after PMA+ Ionomycin stimulation. Data are representative of 2 experiments with 3-5 mice per group. (E) Correlation analysis of the percentage of splenic MMCs versus the percentage of Th17 and Treg cells in female mI Dendra^WT^ offspring. The black line represents the linear regression, and the blue line indicates the nonparametric Spearman correlation with a 95% confidence interval. (F) **Spleen PND28-35:** Frequency of iNKT17 cells, ILC3s and RORγt^+^DCs in offspring of previously infected dams. (mC male n=15, mC female n=16, mI male n=9, mI female n=9; data are from 5 independent experiments using 3-4 pregnant mice dams per group and 4-6 offspring per pregnancy; one-way ANOVA adjusted P: ns not significant). Summary of statistical analyses are shown in **Supplemental Data 1**.

**Supplemental Figure 6:**
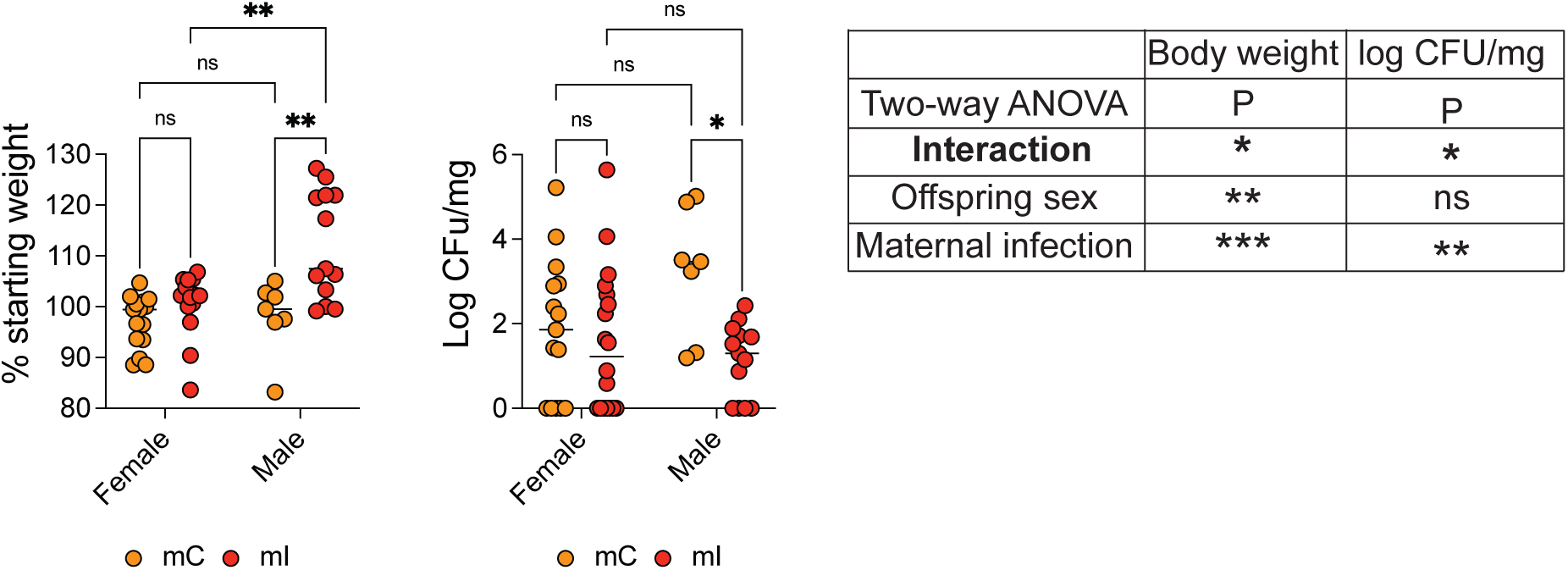
Pregnancy associated infection increases resistance of male offspring to *Salmonella* infection, related to Figure 3 Dendra^WT^ offspring of uninfected (mC) or previously infected dams (mI) were orally administered *Salmonella* Typhimurium (SL1344 mutant) at PND28-35. **(A)** *(Left)* Percent starting weight and (*Right*) CFU 5 days after *Salmonella* infection in offspring of previously infected dams segregated by sex. (mC-Salm M n=7, mC-Salm F n=15, mI-Salm M n=12, mI-Salm F n=18; data are from 4 independent experiments using 3-4 pregnant mice dams per group and 4-6 offspring per pregnancy; two-way ANOVA adjusted P: ns not significant, * <0.033, ** < 0.0021, *** < 0.0002). Summary of statistical analyses are shown in **Supplemental Data 1**.

**Table S1:**
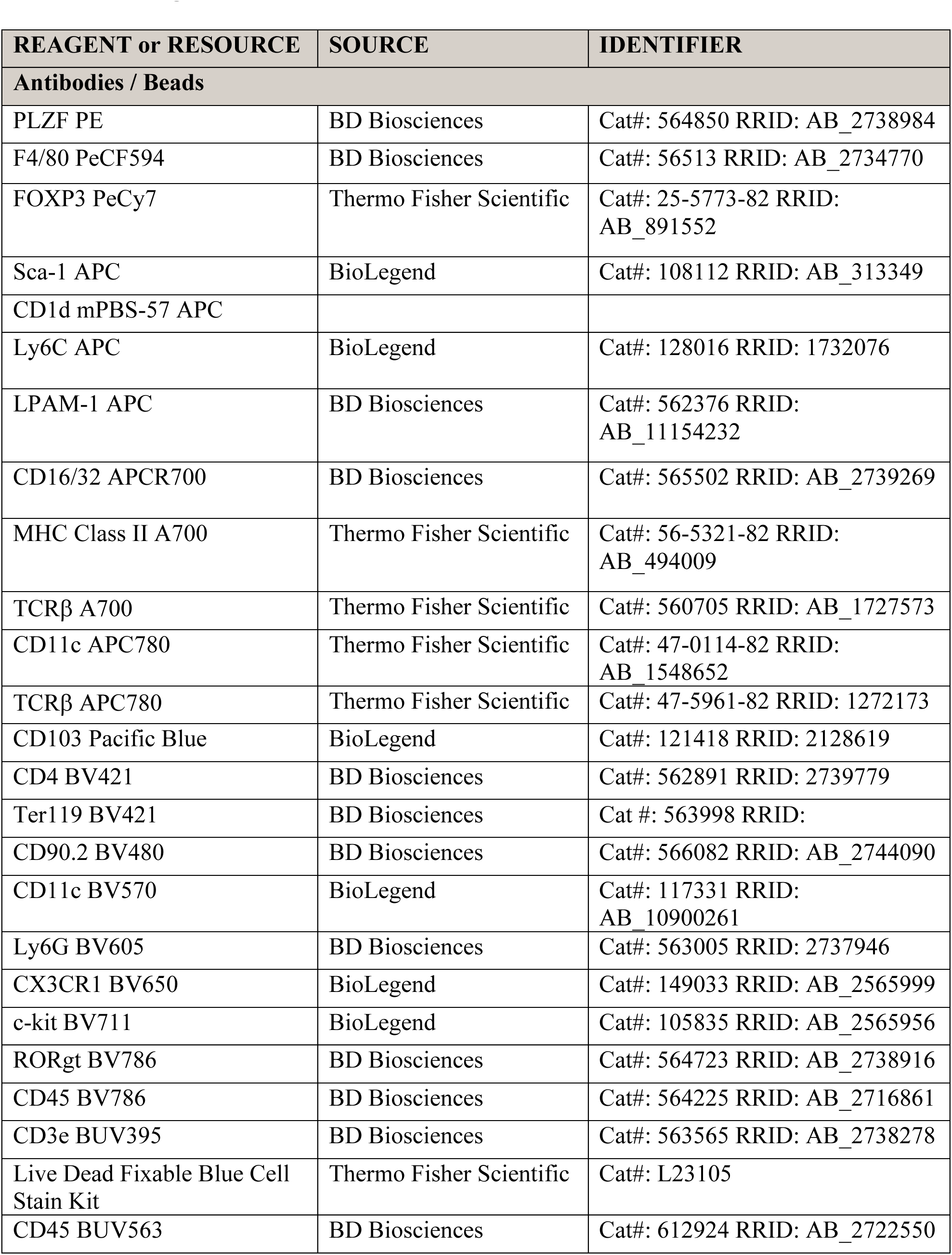
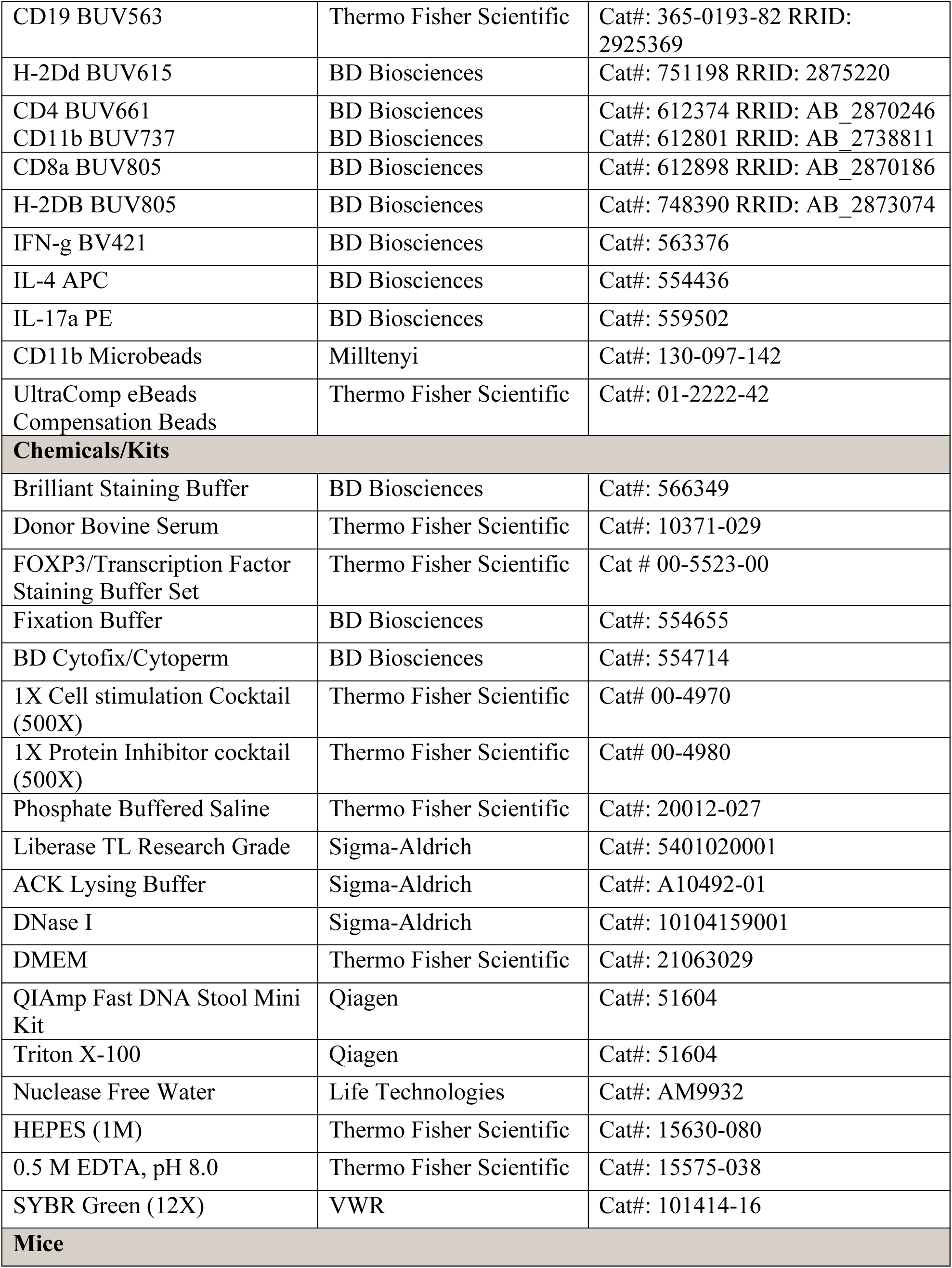
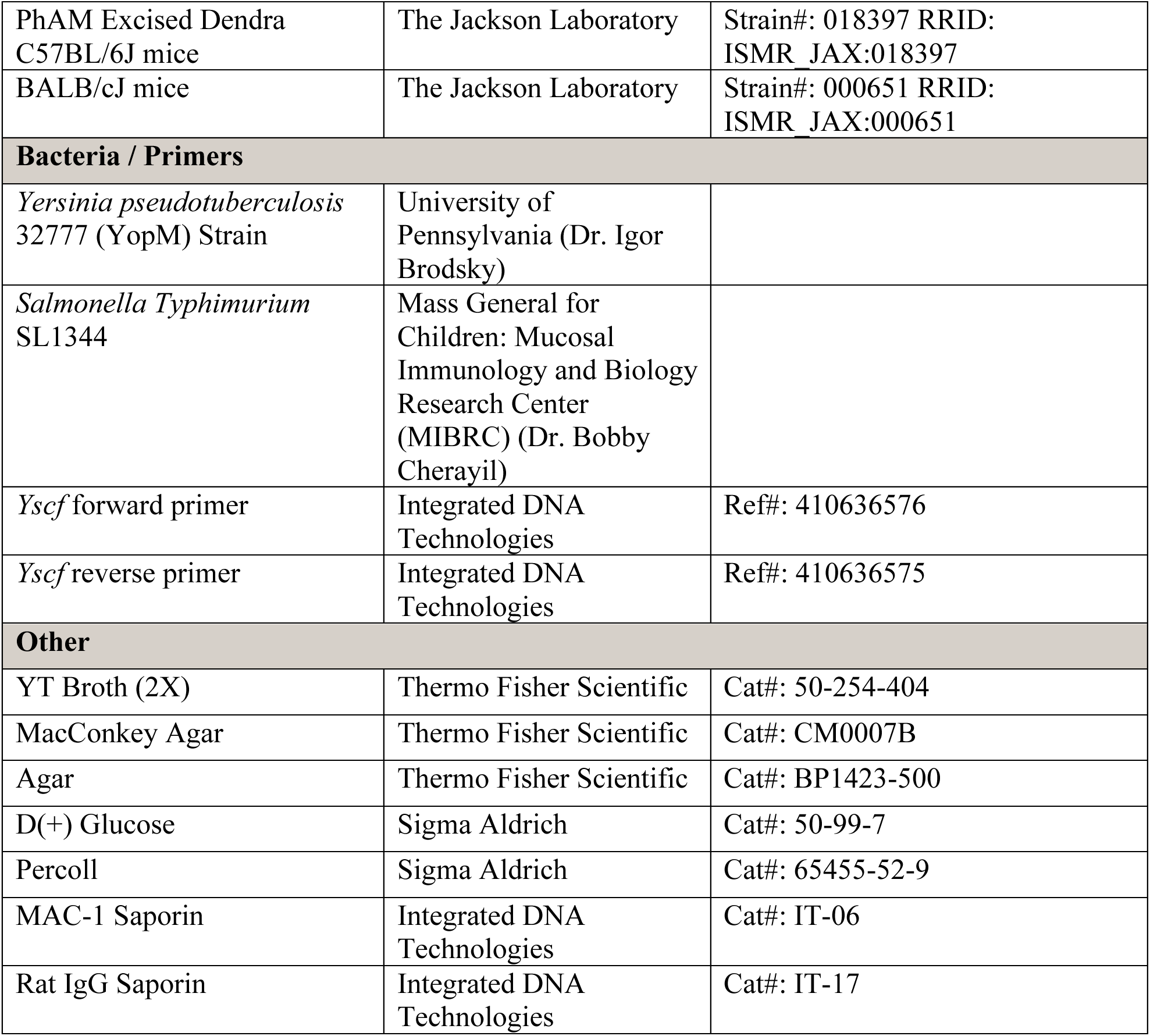
Reagents List.

## Reference

1. N. Dauby, T. Goetghebuer, T. R. Kollmann, J. Levy, A. Marchant, Uninfected but not unaffected: chronic maternal infections during pregnancy, fetal immunity, and susceptibility to postnatal infections. Lancet Infect Dis 12, 330–340 (2012).

2. C. Evans, C. E. Jones, A. J. Prendergast, HIV-exposed, uninfected infants: new global challenges in the era of paediatric HIV elimination. Lancet Infect Dis 16, e92–e107 (2016).

3. I. Malhotra et al., Effect of antenatal parasitic infections on anti-vaccine IgG levels in children: a prospective birth cohort study in Kenya. PLoS Negl Trop Dis 9, e0003466 (2015).

4. J. M. Kinder, I. A. Stelzer, P. C. Arck, S. S. Way, Immunological implications of pregnancy- induced microchimerism. Nat Rev Immunol 17, 483–494 (2017).

5. J. L. Nelson, The otherness of self: microchimerism in health and disease. Trends Immunol. 33, 421–427 (2012).

6. S. Maloney et al., Microchimerism of maternal origin persists into adult life. The Journal of clinical investigation 104, 41–47 (1999).

7. A. M. Stevens, H. M. Hermes, J. C. Rutledge, J. P. Buyon, J. L. Nelson, Myocardial-tissue-specific phenotype of maternal microchimerism in neonatal lupus congenital heart block. Lancet 362, 1617–1623 (2003).

8. E. Roy et al., Specific maternal microchimeric T cells targeting fetal antigens in beta cells predispose to auto-immune diabetes in the child. J Autoimmun 36, 253–262 (2011).

9. J. L. Nelson et al., Maternal microchimerism in peripheral blood in type 1 diabetes and pancreatic islet beta cell microchimerism. Proc Natl Acad Sci U S A 104, 1637–1642 (2007).

10. K. Khosrotehrani et al., Presence of chimeric maternally derived keratinocytes in cutaneous inflammatory diseases of children: the example of pityriasis lichenoides. J Invest Dermatol 126, 345–348 (2006).

11. M. Arvola et al., Immunoglobulin-secreting cells of maternal origin can be detected in B cell-deficient mice. Biology of reproduction 63, 1817–1824 (2000).

12. L. E. Wrenshall, E. T. Stevens, D. R. Smith, J. D. Miller, Maternal microchimerism leads to the presence of interleukin-2 in interleukin-2 knock out mice: implications for the role of interleukin-2 in thymic function. Cellular immunology 245, 80–90 (2007).

13. F. Touzot et al., Massive expansion of maternal T cells in response to EBV infection in a patient with SCID-Xl. Blood 120, 1957–1959 (2012).

14. J. Y. Koh et al., Impact of maternal engrafted cytomegalovirus-specific CD8(+) T cells in a patient with severe combined immunodeficiency. Clin Transl Immunology 10, e1272 (2021).

15. D. Yuzen et al., Pregnancy-induced transfer of pathogen-specific T cells from mother to fetus in mice. EMBO Rep 10.15252/embr.202356829, e56829 (2023).

16. J. E. Mold et al., Maternal alloantigens promote the development of tolerogenic fetal regulatory T cells in utero. Science 322, 1562–1565 (2008).

17. J. M. Kinder et al., Cross-Generational Reproductive Fitness Enforced by Microchimeric Maternal Cells. Cell 162, 505–515 (2015).

18. I. A. Stelzer et al., Vertically transferred maternal immune cells promote neonatal immunity against early life infections. Nat Commun 12, 4706 (2021).

19. C. Balle et al., Factors influencing maternal microchimerism throughout infancy and its impact on infant T cell immunity. The Journal of clinical investigation 132 (2022).

20. E. E. Thompson et al., Maternal microchimerism protects against the development of asthma. The Journal of allergy and clinical immunology 132, 39–44 (2013).

21. K. Shankar et al., Maternal obesity promotes a proinflammatory signature in rat uterus and blastocyst. Endocrinology 152, 4158–4170 (2011).

22. N. V. Malkova, C. Z. Yu, E. Y. Hsiao, M. J. Moore, P. H. Patterson, Maternal immune activation yields offspring displaying mouse versions of the three core symptoms of autism. Brain Behav Immun 26, 607–616 (2012).

23. M. Kumar, M. Saadaoui, S. Al Khodor, Infections and Pregnancy: Effects on Maternal and Child Health. Front Cell Infect Microbiol 12, 873253 (2022).

24. A. H. Pham, J. M. McCaffery, D. C. Chan, Mouse lines with photo-activatable mitochondria to study mitochondrial dynamics. Genesis 50, 833–843 (2012).

25. M. Wegorzewska, T. Le, Q. Tang, T. C. MacKenzie, Increased maternal T cell microchimerism in the allogeneic fetus during LPS-induced preterm labor in mice. Chimerism 5, 68–74 (2014).

26. A. I. Lim et al., Prenatal maternal infection promotes tissue-specific immunity and inflammation in offspring. Science 373 (2021).

27. J. B. McPhee, P. Mena, J. B. Bliska, Delineation of regions of the Yersinia YopM protein required for interaction with the RSK1 and PRK2 host kinases and their requirement for interleukin-10 production and virulence. Infect Immun 78, 3529–3539 (2010).

28. M. Giassi et al., In utero position matters for littermate cell transfer in mice: an additional and confounding source with maternal microchimerism. Front Immunol 14, 1200920 (2023).

29. V. Bronte et al., Recommendations for myeloid-derived suppressor cell nomenclature and characterization standards. Nat Commun 7, 12150 (2016).

30. C. M. Diester, M. L. Banks, G. N. Neigh, S. S. Negus, Experimental design and analysis for consideration of sex as a biological variable. Neuropsychopharmacology 44, 2159–2162 (2019).

31. F. Fransen et al., BALB/c and C57BL/6 Mice Differ in Polyreactive IgA Abundance, which Impacts the Generation of Antigen-Specific IgA and Microbiota Diversity. Immunity 43, 527–540 (2015).

32. C. Ohnmacht et al., MUCOSAL IMMUNOLOGY. The microbiota regulates type 2 immunity through RORgammat(+) T cells. Science 349, 989–993 (2015).

33. E. Sefik et al., MUCOSAL IMMUNOLOGY. Individual intestinal symbionts induce a distinct population of RORgamma(+) regulatory T cells. Science 349, 993–997 (2015).

34. B. S. Kim et al., Generation of RORgammat(+) Antigen-Specific T Regulatory 17 Cells from Foxp3(+) Precursors in Autoimmunity. Cell Rep 21, 195–207 (2017).

35. M. J. McGeachy, S. J. McSorley, Microbial-induced Th17: superhero or supervillain? J Immunol 189, 3285–3291 (2012).

36. F. Ferrini et al., Morphine hyperalgesia gated through microglia-mediated disruption of neuronal Cl(-) homeostasis. Nat Neurosci 16, 183–192 (2013).

37. X. Xiong, Y. Zhang, Y. Wen, Diverse functions of myeloid-derived suppressor cells in autoimmune diseases. Immunol Res 72, 34–49 (2024).

38. H. Wu et al., Arginase-1-dependent promotion of TH17 differentiation and disease progression by MDSCs in systemic lupus erythematosus. Sci Transl Med 8, 331ra340 (2016).

39. M. P. Soares, L. Teixeira, L. F. Moita, Disease tolerance and immunity in host protection against infection. Nat Rev Immunol 17, 83–96 (2017).

40. S. L. Klein, K. L. Flanagan, Sex differences in immune responses. Nat Rev Immunol 16, 626–638 (2016).

41. C. C. Sawyer, Child mortality estimation: estimating sex differences in childhood mortality since the 1970s. PLoS Med 9, e1001287 (2012).

42. J. Y. Cephus et al., Testosterone Attenuates Group 2 Innate Lymphoid Cell-Mediated Airway Inflammation. Cell Rep 21, 2487–2499 (2017).

43. J. P. Hoffmann, J. A. Liu, K. Seddu, S. L. Klein, Sex hormone signaling and regulation of immune function. Immunity 56, 2472–2491 (2023).

44. J. G. Markle et al., Sex differences in the gut microbiome drive hormone-dependent regulation of autoimmunity. Science 339, 1084–1088 (2013).

45. L. Chi et al., Sexual dimorphism in skin immunity is mediated by an androgen-ILC2- dendritic cell axis. Science 384, eadk6200 (2024).

46. M. D. Rudolph et al., Maternal IL-6 during pregnancy can be estimated from newborn brain connectivity and predicts future working memory in offspring. Nat Neurosci 21, 765–772 (2018).

47. J. E. Park et al., A cell atlas of human thymic development defines T cell repertoire formation. Science 367 (2020).

48. M. Ennamorati et al., Intestinal microbes influence development of thymic lymphocytes in early life. Proc Natl Acad Sci U S A 117, 2570–2578 (2020).

49. J. Philpott et al., RXRalpha Regulates the Development of Resident Tissue Macrophages. Immunohorizons 6, 366–372 (2022).

50. M. McCausland, Y. D. Lin, T. Nevers, C. Groves, V. Decman, With great power comes great responsibility: high-dimensional spectral flow cytometry to support clinical trials. Bioanalysis 13, 1597–1616 (2021).

